# Deep learning and alignment of spatially-resolved whole transcriptomes of single cells in the mouse brain with Tangram

**DOI:** 10.1101/2020.08.29.272831

**Authors:** Tommaso Biancalani, Gabriele Scalia, Lorenzo Buffoni, Raghav Avasthi, Ziqing Lu, Aman Sanger, Neriman Tokcan, Charles R. Vanderburg, Asa Segerstolpe, Meng Zhang, Inbal Avraham-Davidi, Sanja Vickovic, Mor Nitzan, Sai Ma, Jason Buenrostro, Nik Bear Brown, Duccio Fanelli, Xiaowei Zhuang, Evan Z. Macosko, Aviv Regev

## Abstract

Charting a biological atlas of an organ, such as the brain, requires us to spatially-resolve whole transcriptomes of single cells, and to relate such cellular features to the histological and anatomical scales. Single-cell and single-nucleus RNA-Seq (sc/snRNA-seq) can map cells comprehensively^5,6^, but relating those to their histological and anatomical positions in the context of an organ’s common coordinate framework remains a major challenge and barrier to the construction of a cell atlas^7–10^. Conversely, Spatial Transcriptomics allows for *in-situ* measurements^11–13^ at the histological level, but at lower spatial resolution and with limited sensitivity. Targeted *in situ* technologies^1–3^ solve both issues, but are limited in gene throughput which impedes profiling of the entire transcriptome. Finally, as samples are collected for profiling, their registration to anatomical atlases often require human supervision, which is a major obstacle to build pipelines at scale. Here, we demonstrate spatial mapping of cells, histology, and anatomy in the somatomotor area and the visual area of the healthy adult mouse brain. We devise Tangram, a method that aligns snRNA-seq data to various forms of spatial data collected from the same brain region, including MERFISH^1^, STARmap^2^, smFISH^3^, and Spatial Transcriptomics^4^ (Visium), as well as histological images and public atlases. Tangram can map any type of sc/snRNA-seq data, including multi-modal data such as SHARE-seq data^5^, which we used to reveal spatial patterns of chromatin accessibility. We equipped Tangram with a deep learning computer vision pipeline, which allows for automatic identification of anatomical annotations on histological images of mouse brain. By doing so, Tangram reconstructs a genome-wide, anatomically-integrated, spatial map of the visual and somatomotor area with ∼30,000 genes at single-cell resolution, revealing spatial gene expression and chromatin accessibility patterning beyond current limitation of *in-situ* technologies.

## INTRODUCTION

An Human Cell Atlas^6–8^, and especially a Brain Cell Atlas, should combine high resolution molecular and histological mapping with anatomical and functional data. Achieving this goal relies critically on our ability to relate different levels of biological organization and data modalities to each other. In this context, molecular data, for example mRNA levels, has the potential to provide a unifying connector, especially if it can be quantified both spatially and at single cell resolution. In addition, such spatial measurements then relate cellular features across scales, including histological and anatomical, by mapping through a Common Coordinate Framework (CFF) of an organ^9^. However, to achieve this ultimate goal, we need to overcome two challenges: obtain molecular profiles at high spatial resolution, and relate those to both histological and anatomical information.

Advances in single cell and spatial genomics^10^ opened the way to high resolution spatial profiles, but each of the currently available technologies only addresses some of the challenge of resolving whole transcriptomes in space at single-cell resolution. On the one hand, single cell and single nucleus RNA-seq (sc/snRNA-seq) profile single cells transcriptome-wide, from which we can recover cell types^11^, gene programs^12,13^, and developmental relations^14,15^, but by necessity loses spatial information. Conversely, spatial technologies resolve transcriptomes in space, but are limited in either gene throughput or spatial resolution. In general, targeted *in situ* technologies (*e*.*g*., ISS^16^, MERFISH^1^, smFISH^3^, osmFISH^17^, STARMap^2^, Targeted ExSeq^18^, seqFISH+^19^) are limited in the number measured genes measured, which must be pre-selected and are typically in the hundreds to thousands range, whereas adding more probes can reduce accuracy for some genes^2^. Spatial Transcriptomics methods (*e*.*g*., Spatial Transcriptomics (ST)^4^ (now available commercially as Visium), Slide-seq^20^, or High Definition Spatial Trascriptomics^21^) spatially barcode entire transcriptomes, but with limited capture rate (and substantial “dropouts”, which increase at higher resolution) and a spatial resolution larger than a single cell, ranging from 50-100 μm for ST to 10 μm for Slide-seq.

Computational methods have previously bridged this gap by combining single cell and spatial measurements^22–25^. With targeted, high resolution, spatial measurements, mapping provides spatial measurements for additional genes based on their (mapped) single cell profiles. These methods can reconstruct key landmark genes by leveraging local alignment in transcriptome space^22–24^, or hypotheses such as continuity in gene expression^25^. However, intrinsically sparse or granularly distributed genes are difficult to predict. For measurements at coarse spatial resolution, computational methods aim to deconvolve these data^20,26^, by either learning a program dictionary^20^ or a probability distribution of the data^26^, to infer a cell type composition within a spatial voxel. However, deconvolution is hindered by spatial “dropouts”, where cells types defined by sparse or dim markers are not correctly detected^27^.

In many cases, only histological data is directly available for the specimens collected as part of single cell atlases, but those can serve as a bridge to pre-existing atlases, with measured *in situ* hybridization (ISH) data, and rich anatomical annotations in the context of a Common Coordinate Framework, as in the case of the Allen Brain Atlas^28^. Using these data should allow relating cellular features (*e*.*g*., gene expression, cell types) to the histological or organ scale, especially in the brain. However, typical methods from computer vision for registration of medical images^29,30^ require human supervision, such as identification of a few corresponding anatomical landmarks in experimental and atlas images. Such supervision, albeit minimal, prevents complete automation. A common strategy to remove supervision uses machine learning for identifying the few key landmarks required in registration, as has been shown for registration of sagittal mouse brain slides^31^. However, this method is not suitable for images that are torn or contain holes, for example, if tissue has been first dissected for profiling sc/snRNA-seq data.

Here, we present Tangram, a deep learning framework to address both challenges: learn spatial gene expression maps transcriptome-wide at single cell resolution, and relate those to histological and anatomical information from the same specimens. Tangram learns a spatial alignment of sc/snRNA-seq data from a reference spatial data of any kind – either fine or coarse grained, as we demonstrate by spatially mapping snRNA-seq data from the isocortex of the adult healthy mouse brain using each of five kinds of spatial supports, at different levels of resolution and gene coverage: ISH, smFISH, Visium (Spatial Transcriptomics), STARmap and MERFISH. Tangram produces consistent spatial maps of cell types, overcomes limitations in throughput or resolution, corrects low-quality genes, even in high resolution methods, provides single cell resolution for low-resolution methods, and provides genome-wide coverage for targeted methods. By mapping multi-modal single data (SHARE-seq^5^) on spatial support, Tangram visualizes spatial patterns of chromatin accessibility and transcription factor motif scores at single cell resolution. Finally, Tangram includes a dedicated novel computer vision module that leverages histological data, and maps it to anatomical positions in an existing Common Coordinate Framework in the brain. If a histology image is available, even without any further annotation, this module relates all scales, to a single integrated atlas.

## RESULTS

### Tangram: Learning of spatially-resolved single cell transcriptomes by alignment

We developed Tangram, an algorithm that uses sc/snRNA-seq as “puzzle pieces” to align in space to match “the shape” of the spatial data (**Fig. 1a**). The input to Tangram is sc/snRNA-seq data along with spatial data from the same region or tissue type, from any currently available spatial method (*e*.*g*., MERFISH, smFISH, STARmap, ISH, or Visium), requiring only that the two modalities share at least some subset of common genes. Intuitively, Tangram first randomly places the sc/snRNA-seq profiles in space, then computes an objective function that mimics the spatial correlation between each gene in the sc/snRNA-seq data and in the spatial data. Tangram then rearranges the sc/snRNA-seq profiles in space to maximize the total spatial correlation across the genes shared by the datasets. When Tangram finishes, the mapped sc/snRNA-seq profiles constitute the new spatial data: they now contain all genes at single cell resolution and with spatial position. From the learned mapping function, Tangram can (**1**) expand from a measured subset of genes to genome wide profiles (**Fig. 1b**); (**2**) correct low-quality spatial measurements (**Fig. 1c**); (**3**) map the location of cells of different types (**Fig. 1d**); (**4**) deconvolve low resolution measurements to single cells (**Fig. 1e**); and (**5**) resolve spatial patterns of chromatin accessibility at single-cell resolution by aligning multi-modal data (**Fig. 1e**).

**Figure 1.**
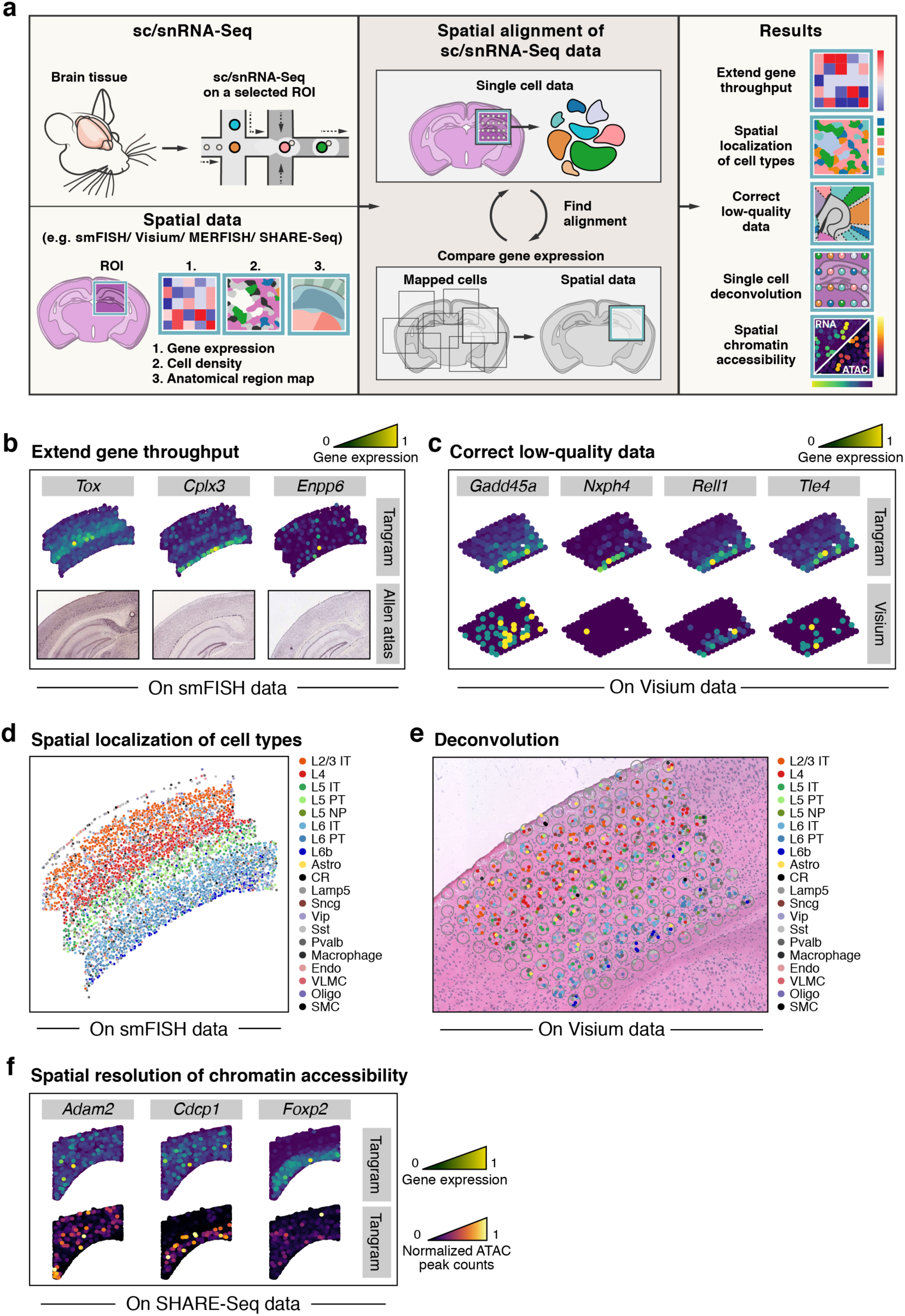
Tangram learns spatial transcriptome-wide patterns at single cell resolution from sc/snRNA-seq data and corresponding spatial data. **a**. Overview. sc/snRNA-seq data and spatial data, collected from the same tissue are spatially aligned by comparing gene expression of their shared genes. **b-f**. Tangram use cases. **b**. Generating genome wide spatial patterns from gene signature data. Predicted expression patterns (color bar, normalized mRNA counts, **Methods**) for each of three genes not included in an input smFISH dataset validated against their corresponding images from the Allen ISH atlas (bottom). **c**. Correction of low-quality data for spatially measured genes. Predicted (top) and measured (bottom, by Visium) expression patterns (color bar, normalized mRNA counts, **Methods**) of four known markers whose correct localization is missing in direct Visium measurements, but recovered in the predicted patterns. **d**. Cell type localization. Spatial distribution of cell types defined by snRNA-seq (legend) mapped on a an smFISH brain slide. **e**. Single cell deconvolution of lower-resolution Spatial Transcriptomics. Predicted single cells (colored dots, legend) in each Visium voxel (grey circle) based on snRNA-seq data mapped onto a Visium slide. **f**. Spatially resolved chromatin patterns. Predicted spatial gene expression (top, color bar, normalized mRNA counts, **Methods**) and chromatin accessibility (bottom, color bar, normalized ATAC peak counts, **Methods**) by mapping the RNA component of SHARE-Seq data to a MERFISH slide.

Technically, Tangram is based on non-convex optimization (for full details, see **Methods**), where the Tangram optimization function rewards the spatial alignment of sc/snRNA-seq data using two similarity functions: cell density distributions are compared using Kullback-Leibler (KL) divergence, whereas gene expression is assessed via cosine similarity. If the sc/snRNA-seq data contains more cells than the spatial data (which is the typical case), a filter term in the loss function ensures that only the optimal subset of sc/snRNA-seq observations is selected. The output of Tangram is a probabilistic mapping, namely, a matrix denoting the probability of finding each cell from the sc/snRNA-seq data in each voxel of the spatial data. From this matrix, we can obtain a deterministic mapping by assigning the most likely sc/snRNA-seq cell to each spatial voxel. Optionally, we can also activate an entropy regularizer to ensure that the spatial probabilities of each cell are peaked over a narrow portion of space. We did not need to use such feature, as all probabilities were peaked in practice in all cases analyzed in this study.) Tangram does not contain any hyperparameters, maps a hundred thousand cells in a few minutes (using a single P100 GPU), and is released as PyTorch module.

### Tangram maps cells with MERFISH measurements to generate genome-scale high resolution expression maps

To apply Tangram, we collected 160,000 snRNA-seq profiles using droplet-based RNA-seq (10Xv3, see for example^32^), as part of the BRAIN Initiative Cell Census Network (BICCN), from the primary motor area (MOp) of health adult mouse brain. Each profile contains the expression of ∼27,000 genes, and was annotated following the recently delineated cell type taxonomy of neocortical areas^33^, to 22 subclasses (hereafter, “cell types”)^34^. We mapped these snRNA-seq data with a high resolution MERFISH dataset of 254 genes, on a section segmented to 4,234 cells (**Fig. 2**). We trained Tangram using 253 MERFISH genes (all genes but one were detected in our snRNA-seq data). 50% of the aligned profiles were neuronal, with a 6:1 ratio between glutamatergic and GABAergic cells, in accordance with their ratios in snRNA-seq.

**Figure 2.**
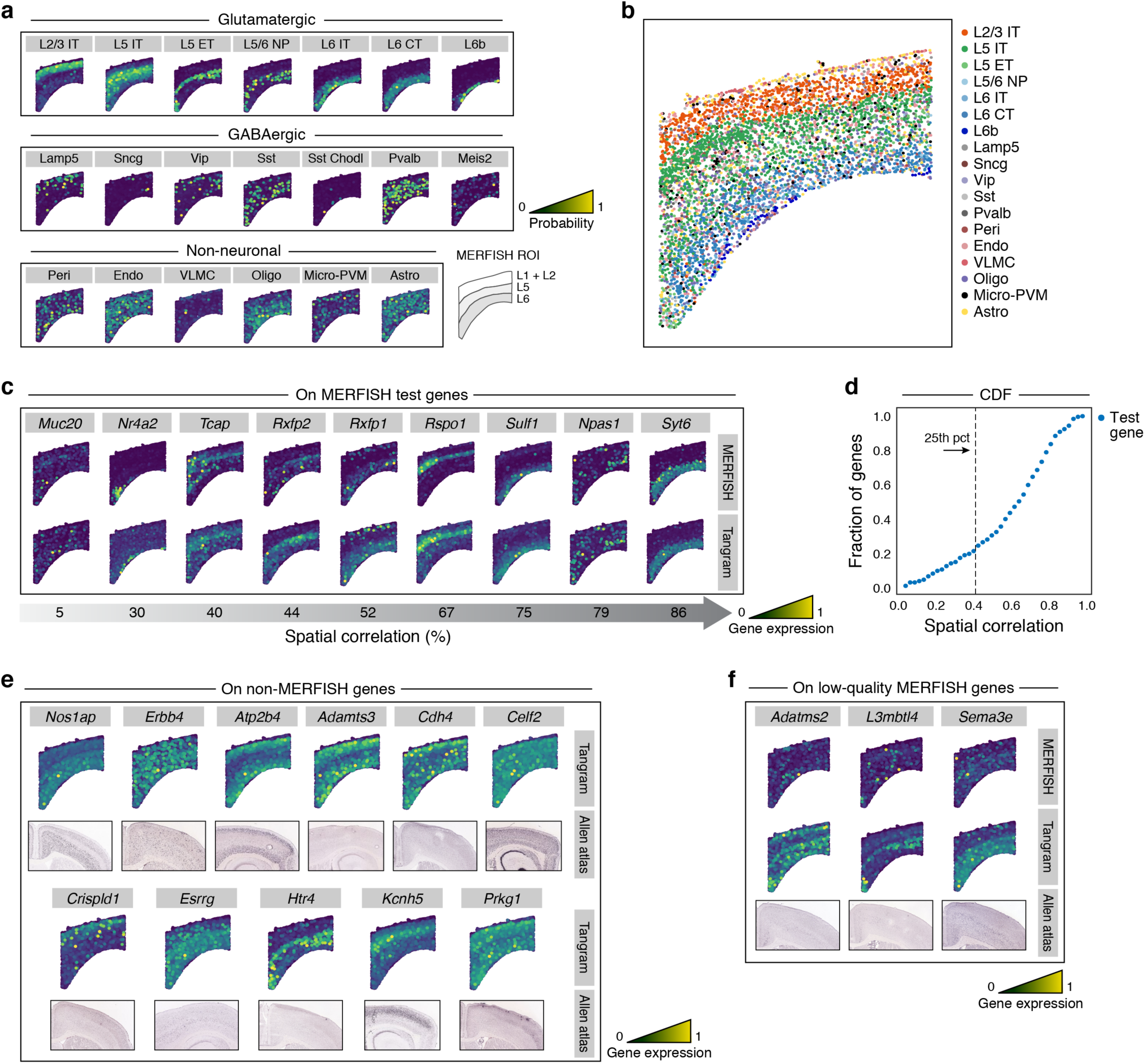
Tangram maps cells with high resolution MERFISH measurements and expands them to genome scale. **a**. Probabilistic mapping of snRNA-seq data on MERFISH data. Probability of mapping (color bar) of each cell subset (grey label) in each of 3 major categories. Bottom right: schematic of key layers. **b**. Deterministic mapping. MERFISH slide with segmented cells (dot) colored by the cell type annotation of the most likely snRNA-seq profile mapped on that position by tangram (legend). **c**,**d**. Predicted expression of test genes. **c**. Measured (top) and Tangram-predicted (bottom) expression (color bar signifies fluorescence at top and normalized mRNA counts at bottom, **Methods**) of select test gene (grey labels) with different extents of spatial correlation (bottom arrow, %) between measured and predicted patterns. **d**. Cumulative distribution function (CDF) of spatial correlation (*x* axis) between predicted and measured patterns for test genes. Dashed line: 75% of test genes are predicted with spatial correlation >40%. **e**. Predicted expression of test genes. Tangram-predicted (bottom) expression (top; color bar, normalized mRNA counts, **Methods**) and corresponding ISH images from the Allen Brain Atlas (bottom) for 11 genes not measured by MERFISH. **f**. Correction of low-quality spatial measurements. MERIFSH measured (top), Tangram-predicted (middle) and Allen Brain Atlas ISH, for genes whose predicted patterns differ from MERFISH measurement but match direct inspection of Allen ISH images (color bar, normalized mRNA counts, **Methods**).

To reveal the spatial distribution of cell types, we combined the learned probabilistic mapping with the cell type annotations in the snRNA-seq data, and obtained the spatial probability distribution for each cell type (**Fig. 2a**). Glutamatergic cells show distinct cortical layer patterns of neuronal subpopulations whereas most, but not all, non-neuronal cells and GABAergic neurons are granularly distributed, as expected. Exceptions include non-neuronal *VLMC* cells (strongly localized in the first layer) and GABAergic *Vip* and *Lamp5* cells, which appear more concentrated toward the upper layers. To verify that these distributions were not an artifact of our probabilistic approach, we also visualized the cell type assignment from the deterministic mapping (*i*.*e*. only the most-likely cell is assigned to each spatial location) and observed similar patterns (**Fig. 2b**).

The learned Tangram model could predict spatial expression patterns well, as we demonstrated by a leave-one-out analysis. Specifically, we partitioned the genes into 252 training genes and a single test gene (unseen in the learning of the model). We repeated the training 253 times, each time leaving out a different gene, to obtain prediction for each gene. As an evaluation score, we computed the spatial correlation between each gene’s real measurement and the predicted spatial pattern of that gene by the learned model. Overall, 75% of the 253 MERFISH genes are predicted with correlation > 40% (**Fig. 2d**). To interpret these spatial correlations, we chose nine genes with varied scores and visually compared the predicted spatial patterns to the MERFISH measurements (**Fig. 2c**). Importantly, the spatial patterns had good qualitative agreement for a broad range of spatial correlation values. For example, the prediction for *Tcap* (40% correlation) is in good accordance with its measurement. This is because when spatial resolution is at the single-cell level, correlation is highly sensitive to noise in gene expression or its measurement, such that somewhat lower correlation does not imply qualitative disagreement. This phenomenon is especially evident in highly sparse genes (*e*.*g*., *Muc20*): the sparse pattern is recapitulated, but the specific single cell locations are not precise, which may reflect the true nature of these patterns.

Mapping snRNA-seq data on MERFISH increases gene throughput to 27,000 genes, which we validated for 11 selected genes with available ISH data in the Allen ISH dataset (**Fig. 2e**). Some genes exhibit strong, localized, patterns in striking similarity to those in the Allen images (*Kcnh5, Nos1ap, Erbb4, Atp2b4, Celf2, Crispld1*). For other genes the signal in the Allen ISH image is very dim compared to our predictions (*Esrrg, Cdh4, Adamts3, Htr4, Prkg1*), but a close inspection reveals agreement as well. This suggests that Tangram can reveal, spatial patterns for lowly-expressed genes, as we will further demonstrate below (with Spatial Transcriptomics Visium data). Interestingly, we obtained similar results when we predicted withheld genes that were measured by MERFISH but had relatively lower-quality, possibly because of less optimal oligonucleotide probes used for these genes: the model predictions were consistent with ISH data, suggesting that the model can “correct” lower-quality signal (**Fig. 2f**). Correction of lower-quality measurements can also partly account for lower correlation scores between predictions and measurements (**Figs. 2c,f**).

### Accurate correction of low-quality transcripts in STARMap

To further investigate Tangram’s correction of low-quality transcripts, we analyzed a STARmap dataset^2^. STARmap is a targeted *in-situ* technology which has been used to measured ∼1,000 genes in 3D resolution; however, increasing the number of measured genes trades-off with gene expression accuracy. In this dataset^2^, 1020 genes are measured in 972 cells in a mouse brain slice from the visual area (VISp). We map 11,759 SMART-Seq2^35^ snRNA-seq profiles from the VISp area using 995 training genes present in both STARmap and snRNA-seq data.

Inspecting cell type distributions from either probabilistic (**Fig. 3a**) or deterministic (**Fig. 3b**) mapping, we confirmed that cell type patterns are consistent with those obtained with MERFISH from the motor area (**Fig. 2a,b**). Despite minor cell-type annotation difference between the VISp and MOp snRNA-seq datasets, our model provides robust mapping. For example, while only the VISp (but not MOp) snRNA-seq dataset has an annotated glutamatergic L4 (layer four) cell subset, the model successfully revealed L4 in the MOp data (**Fig. 3a**) from predicting its marker genes (e.g. *Kcnh5* in **Fig. 3e** and **Fig. 2e**). Finally, the STARmap dataset also contains sub-cortical tissue (defined as cells below the L6b layer), which allows us to further validate predictions by observing an expected subcortical concentration of oligodendrocytes (**Fig. 3a**).

**Figure 3.**
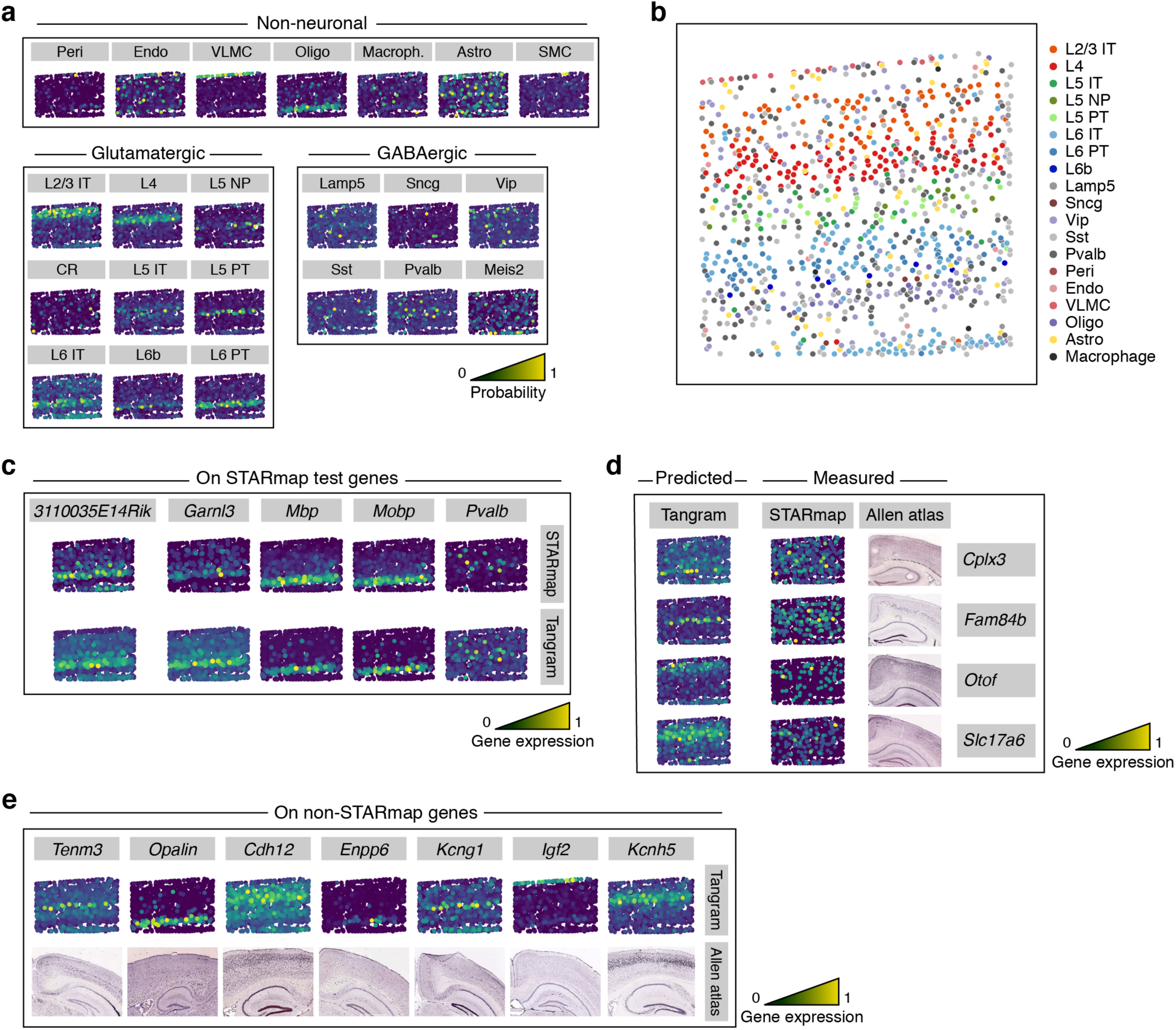
Correction of low-quality genes by mapping snRNA-seq on STARmap data. **a**. Probabilistic mapping of snRNA-seq data on STARmap data. Probability of mapping (color bar) of each cell subset (grey label) in each of 3 major categories. **b**. Deterministic mapping. STARmap slide with segmented cells (dot) colored by the cell type annotation of the most likely snRNA-seq profile mapped on that position by Tangram (legend). **c**. Measured (top) and Tangram-predicted (bottom) expression (color bar signifies fluorescence at top and normalized mRNA counts at bottom, **Methods**) of select test gene (grey labels). **d**. Correction of low-quality spatial measurements. Tangram-predicted test genes (left), STARmap measurements (middle) and Allen atlas images (right) (color bar, normalized mRNA counts, **Methods**) of four genes (grey labels) whose predicted patterns differ from STARmap measurement but match direct measurement by MERFISH. **e**. Predicted expression of test genes. Tangram-predicted (top) expression (top; color bar, normalized mRNA counts, **Methods**) and corresponding ISH images from the Allen Brain Atlas (bottom) for 6 genes not measured by STARmap.

Remarkably, Tangram not only predict expression for genes that were not measured by STARMap, but effectively corrects the spatial expression of low-quality genes (**Fig. 3c,d,e**), as compared to Allen Brain Atlas ISH. First, when holding out each individual STARmap gene from the training, the predicted expression was typically consistent with direct measurements (**Fig. 3c**). Interestingly, for some genes our predicted localized patterns were not observed in measurements, especially for lower-quality genes (**Fig. 3d**). Remarkably, in these cases, the predicted pattern agreed well with images from the Allen Brain ISH Atlas (**Fig. 3d**), confirming the accuracy of our predictions, and Tangram’s ability to correct gene expression of low-quality data. Finally, Tangram correctly predicted the expression of genes that were not measured by STARMap, including markers of cortical layers (*Tenm3, Cdh12, Kcng1, Igf2*) or sub-cortical tissue (*Opalin* and *Enpp6*) as assessed by their consistency with the Allen Brain ISH Atlas (**Fig. 3e**).

### Single cell deconvolution and histological data incorporation with Spatial Transcriptomics

Next, we focused on the deconvolution challenge in the context of lower-resolution Spatial Transcriptomics (Visium) data measuring 31,053 genes within 50 *µm* diameter circular spots in three coronal mouse brain slices (**Fig. 4**). This is followed by an H&E stain of the slice (*section 1*), with ∼160 circular spots on a region of interest. As single cells are visible in the stained images, we segmented cells to directly estimate cell number within each spot, and overall counted 939 cells.

**Figure 4.**
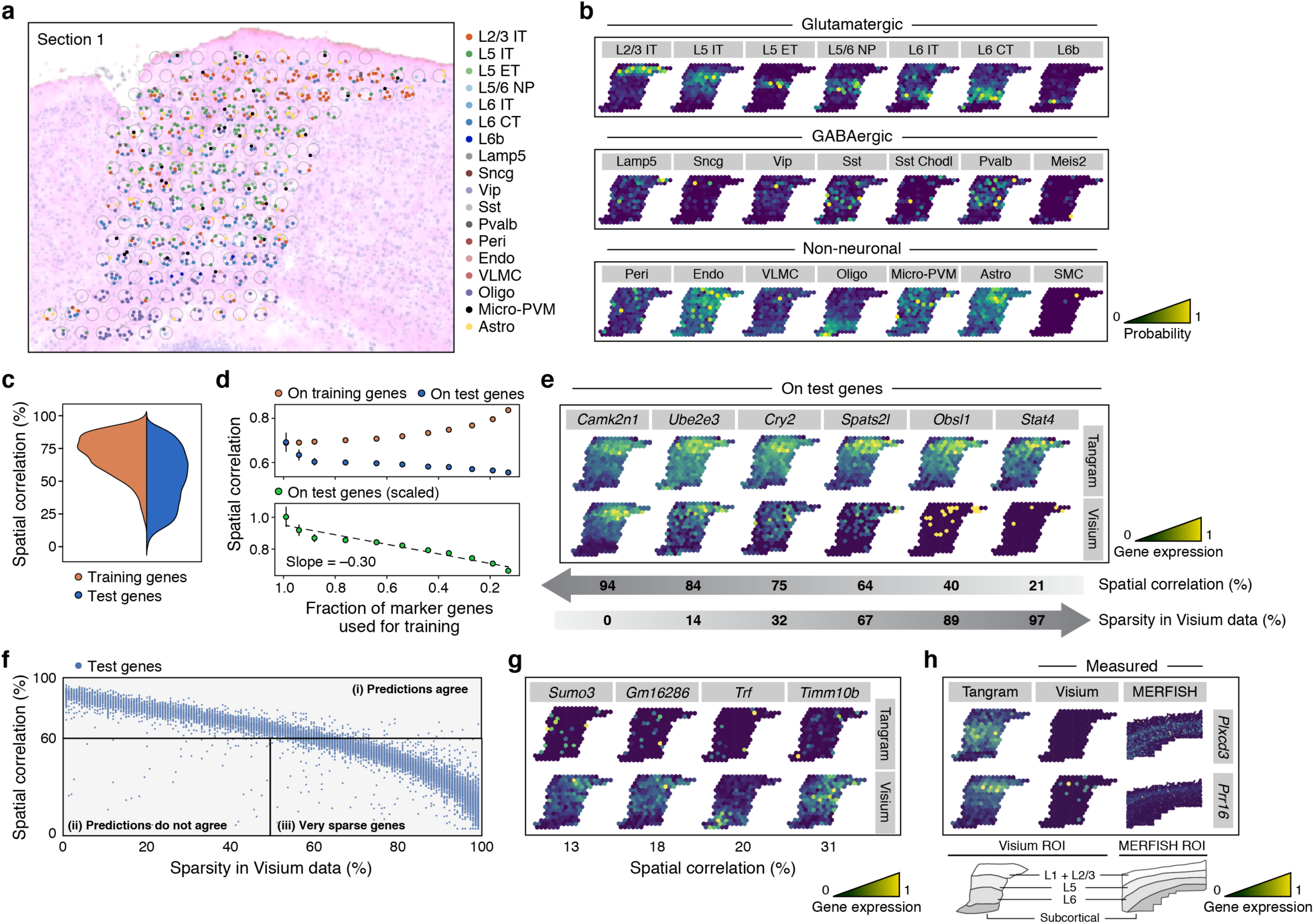
Mapping snRNA-seq data to Spatial Transcriptomics data (Visium) demonstrates deconvolution and imputation of dropouts. **a**. Single cell deconvolution. Predicted single cells (colored dots, legend) in each Visium voxel (grey circle) based on snRNA-seq data mapped onto Visium slide. Cell assignment within a voxel is random with respect to the specific segmented cell. **b**. Probabilistic mapping of snRNA-seq data on the Visium ROI. Probability of mapping (color bar) of each cell subset (grey label) in each of 3 major categories. **c,d**. Predicted expression of test and training genes. **c**. Normalized (i.e. unit area) distribution of single-gene spatial correlation coefficients (<u>y</u> axis) between Tangram-predicted and Visium measured patterns in training (orange) and test (blue) genes. **d**. Reducing the number of training genes decreases prediction performance. Spatial correlation (*y* axis, top) for training gene (orange) and test genes (blue), and scaled spatial correlation (y axis, bottom) for test genes (scaled by the correlation averaged across training genes) for Tangram models learned with different fractions of 1,237 input training genes (*x* axis). **e-h**. Impact of Visium data sparsity on prediction and correction. **e**. Tangram-predicted (top) and Visium measured (bottom) expression (color bar, normalized mRNA counts, **Methods**) of six select test genes (grey labels) with different extents of spatial correlation between measured and predicted patterns (top arrow, %) and of Visium data sparsity (bottom arrow, %). **f**. Spatial correlation of test genes is negatively correlated to sparsity in Visium data. Spatial correlation (*y* axis) between measured and predicted patterns for test genes (blue dots) and their corresponding measurement sparsity (*x* axis). Lines delineate three regions according to model performance. **g**. Few low sparsity genes are not predicted well. Tangram-predicted (top) and Visium measured (bottom) expression (color bar, normalized mRNA counts, **Methods**) of four genes (grey labels) with low sparsity that are not well-predicted by model (from region (ii) of panel f). **h**. Correction of low-quality spatial measurements. Tangram-predicted (left), Visium (middle) and MERFISH (right) measurements (color bar signifies fluorescence for MERFISH figure, normalized mRNA counts for all others, **Methods**), of two genes (grey labels) whose predicted patterns differ from Visium measurements but match direct measurement by MERFISH, and are highly-sparse in Visium measurements (from region (iii) of panel f).

In Spatial Transcriptomics, unlike MERFISH and STARMap, we possess a histological image, which identifies a specific depth in the brain coronal sectioning (more precisely, an anterior-to-posterior coordinate). This opens up two strategies for mapping: (1) map the full MOp snRNA-seq data, constraining the cell number to 939 (as for MERFISH and STARmap); this is more suitable for deconvolution, given the direct assignment on segmented cells; alternatively (2) we can leverage the known anterior-to-posterior coordinate, and only map MOp snRNA-seq profiles collected at that depth (with a method we describe later on); this should resolve smoother gene patterns, given the higher number of profiles as we show in the following section.

For deconvolution, we performed a deterministic mapping of each of the cells within each 50 *µm* voxel, to obtain cell type localization prediction at single-cell resolution (**Fig. 4a**). We trained Tangram with a subset of the >30,000 genes, by selecting 1,237 training genes as the union of the top 100 marker genes of each cell type in the MOp snRNA-seq data (using a standard pipeline^20^, **Methods**) that are detected in the Visium profiles. Unlike probabilistic mapping, we assigned a discrete number of cells to each voxel (matching the number of segmented cells). Notably, rare cell types might not be assigned by deterministic mapping (*e*.*g. Meis* GABAergic neurons).

Our deterministic mapping was successful, as demonstrated by comparing the mapped cell type ratios and those from the snRNA-seq data, which were consistent within Poissonian error (**Extended Data Fig. 1b**). Our mappings were also robust, as demonstrated by analyzing two other Visium datasets: a coronal section (*section 2*) consecutive to *section 1* from the same specimen, and a coronal section collected at approximately the same posterior, provided publicly^36^ (*section 3*) (**Extended Data Fig. 1c**). We note that assignment within a voxel is random: *e*.*g*., the model may predict that one microglia cell is contained in a certain voxel but not *which* cell it is.

### Tangram imputation of dropouts in Spatial Transcriptomics

Next, leveraging the known anterior-to-posterior coordinates, we mapped only MOp snRNA-seq profiles collected at that depth to impute gene expression and correct low-quality data (mostly due to dropouts). Specifically, during collection our MOp snRNA-seq data, we tracked the dissected region, which we annotated as “anterior”, “mid” or “posterior” based on its own histological data, as we describe below (**Fig. 7a, Methods**). As all three Visium histological images were called as “posterior” MOp, we only mapped the 58,002 snRNA-seq profiles from “posterior” samples, using probabilistic mapping.

**Figure 5.**
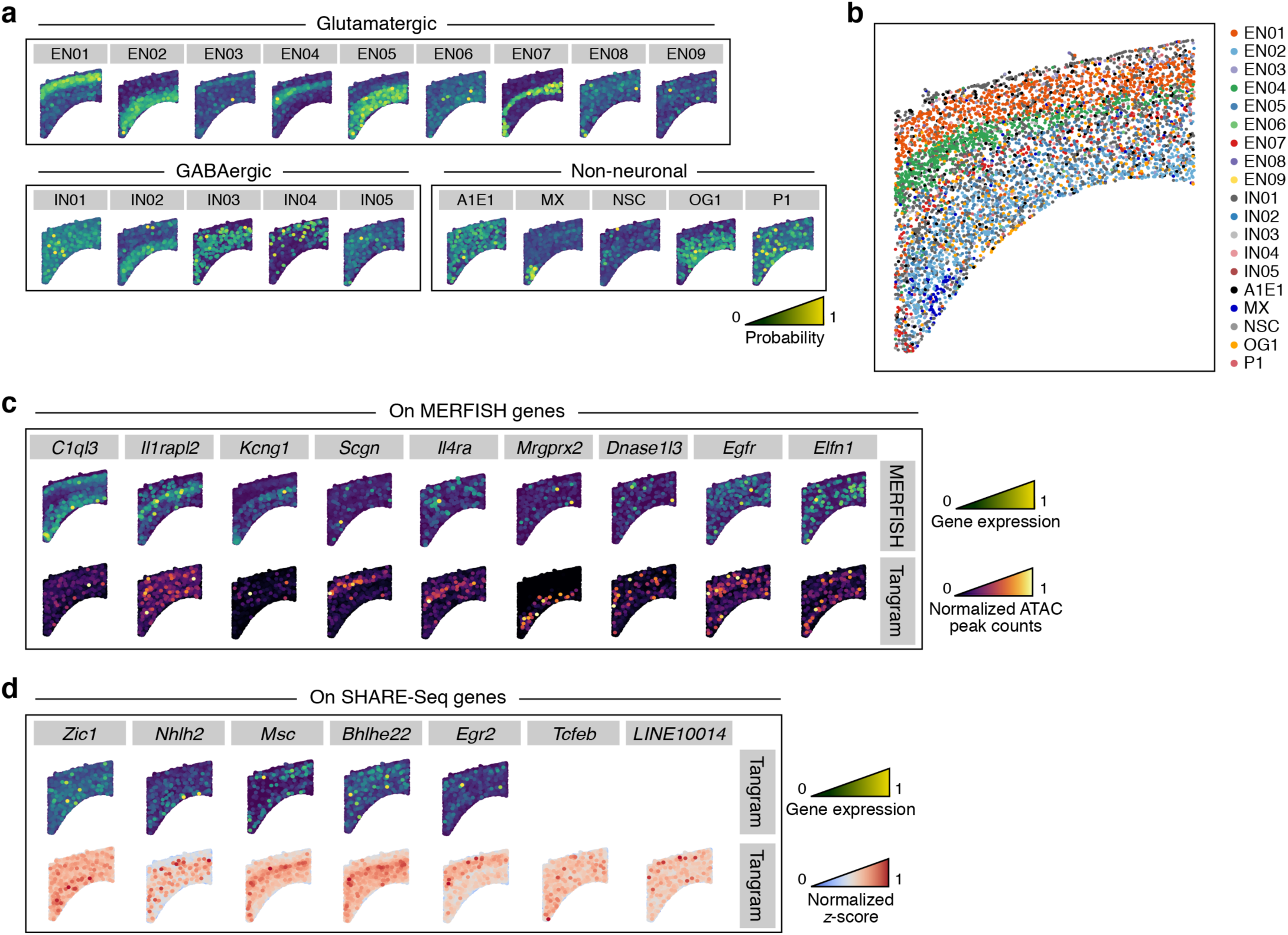
Tangram mapping of multi-omic SHARE-Seq profiles yields spatial patterns of chromatin accessibility and TF activity. **a**. Probabilistic mapping of SHARE-seq profiles on MERFISH data. Probability of mapping (color bar) of each cell subset (grey label) in each of 3 major categories based on the RNA component of SHARE-Seq profiles. **b**. Deterministic mapping. MERFISH slide with segmented cells (dot) colored by the cell type annotation of the most likely SHARE-Seq (RNA) profile mapped on that position by Tangram (legend). **c**. Predicted chromatin accessibility patterns. MERFISH measured expression (top, color bar, normalized fluorescence, **Methods**) and Tangram-predicted chromatin accessibility (bottom, color bar, normalized reads-in-peak count, **Methods**) of select genes (grey labels). **c**. Predicted TF activity patterns. Tangram predicted expression (top, color bar, mRNA counts) and activity normalized z-scores patterns (as inferred from snATAC-Seq, see **Methods**) (bottom, color bar, dimensionless) of select genes (grey labels) measured only by SHARE-Seq.

**Figure 6.**
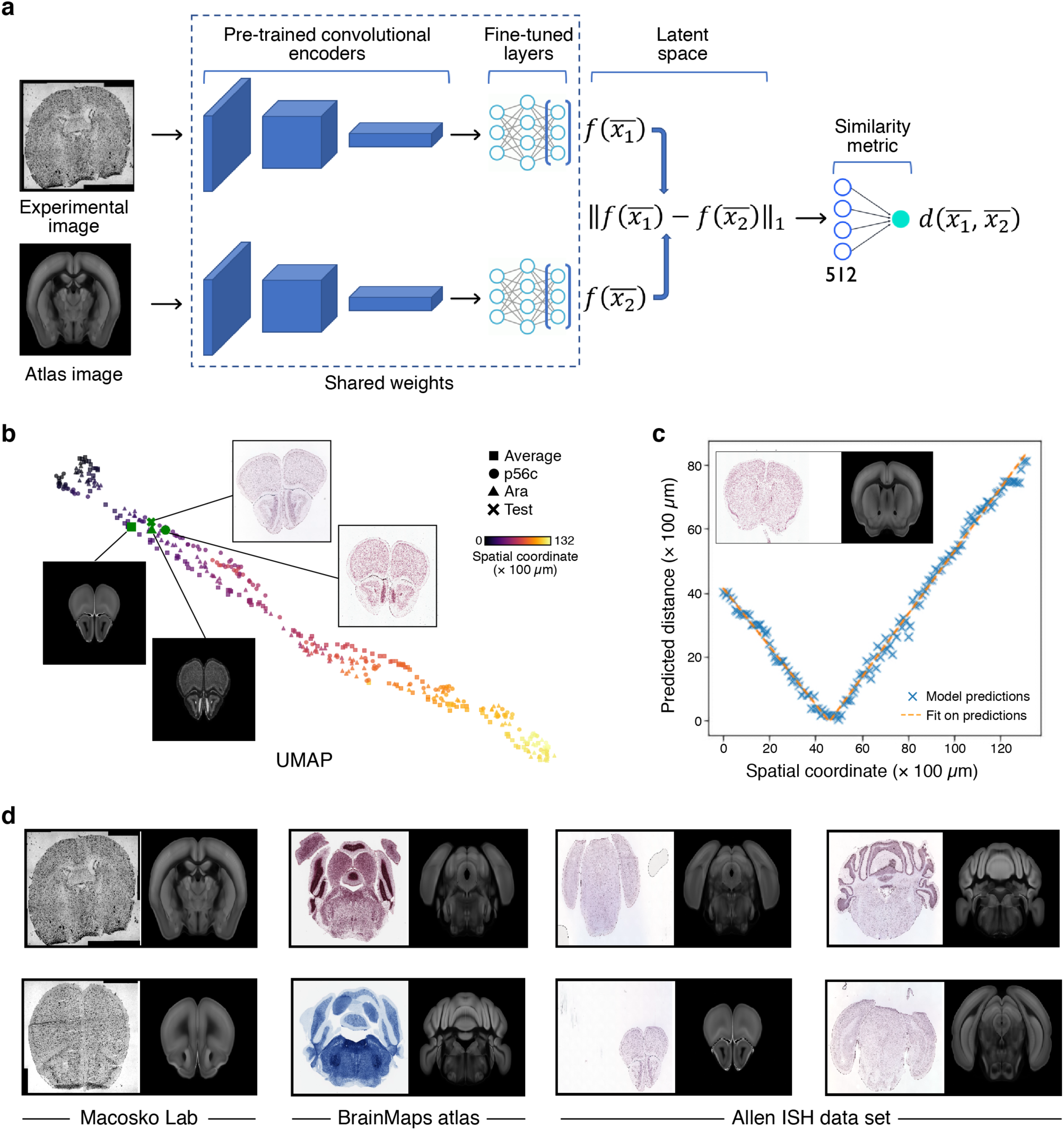
A Siamese network model learns a similarity metric for brain sections based on anatomical landmarks in mouse brain images. **a**. Schematic of neural network architecture. A pair of images is fed to two convolutional encoders, which encode them into a 512-dimensional latent space. The image pair is labeled by the spatial coordinate (*i*.*e*., coronal depth) difference between the two images. **b**. The learned latent space is a 1D-manifold ordered by spatial coordinates. UMAP plot of the encoded training images from individual atlases (legend) colored by spatial depth (color bar). Insets illustrate four anatomically similar images from three different atlases and a test image. **c**. Prediction of spatial coordinates for a test image. **c**. Predicted spatial coordinate distance (*y* axis) between a test image (inset, left panel) and each image of the training set obtained at different spatial coordinates (*x* axis). Dashed orange line: |*ax* + *b*| fit via mean square error minimization (*a*∼ − 0.96, *b*∼43).The minimum of the fit is the predicted spatial coordinate (associated image is in the inset, right panel). **d**. Examples of model predictions (right) on test images (left) from the Macosko lab (first column; **Methods**), BrainMaps atlas (second column) and Allen ISH dataset (third and fourth columns).

**Figure 7.**
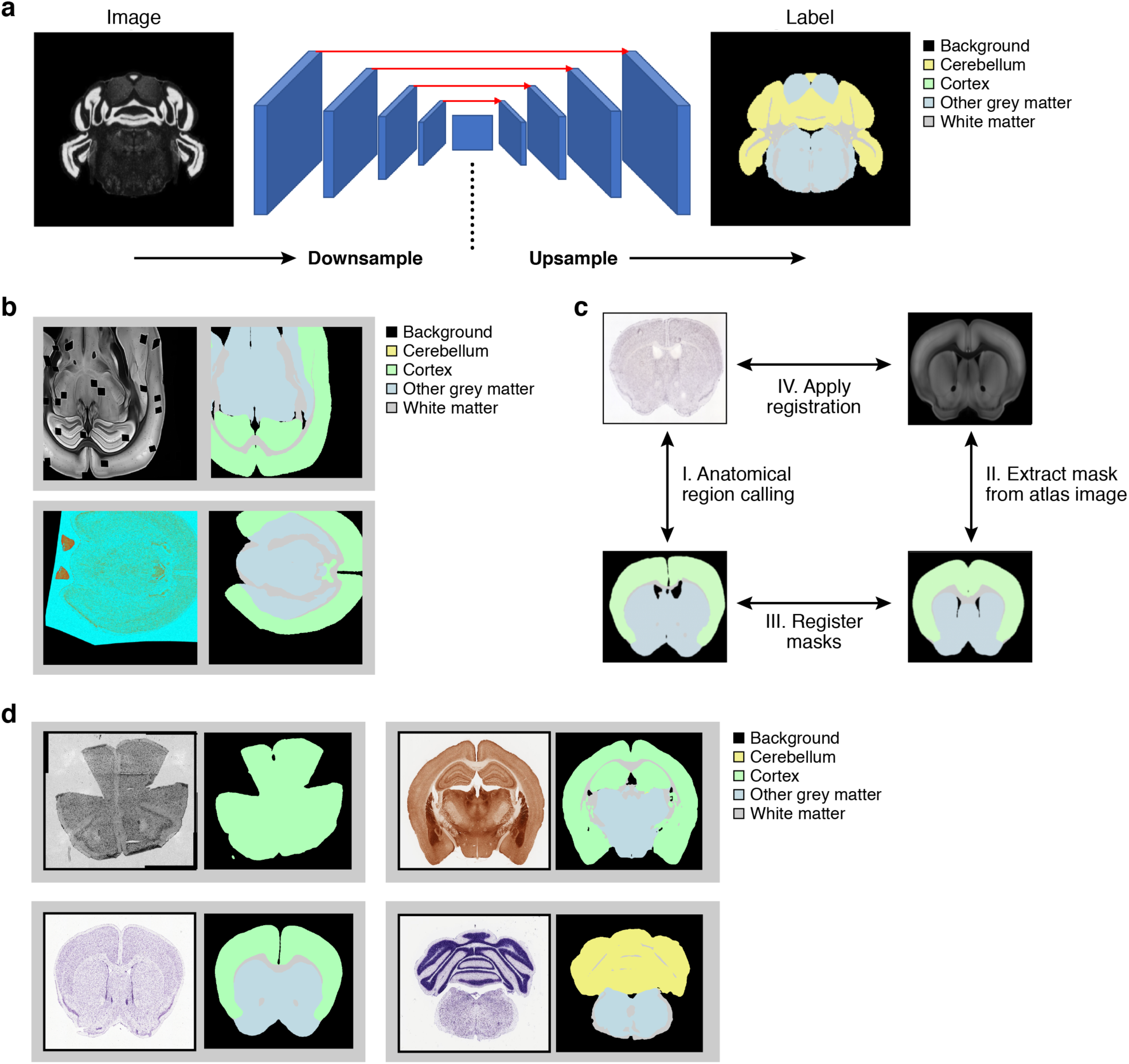
Anatomical region calling via semantic segmentation. **a**. Neural network model used for semantic segmentation. A U-net model is trained on mouse brain images from Allen atlas (left) to recognize five different classes on a mouse brain image (color legend, right). **b**. Augmentation pipeline. Each image undergoes a series of stochastic transformations including affine displacements, dropouts and color shifts (**Methods**). Four training samples are shown. **c**. Schematic of registration strategy. A segmentation mask of an experimental image is produced (I), the mask of each atlas image is extracted in parallel (II), the two masks are registered to each other (III); and finally the learned transformation is used to register the original images (IV). **d**. Prediction examples. Test images (left) and their predicted anatomical region calls (right).

Tangram’s mapping yielded higher resolution, finely localized, cell types (**Fig. 4b, Extended Data Fig. 1a**). This included correct localization of *L6b* glutamatergic neurons, more concentrated presence of *Vip*^*+*^ and *Lamp5*^*+*^ GABAergic neurons in upper layers, positioning of *Sst* and *Pvalb* GABAergic neurons in deeper layers, and of *Meis2*^*+*^ and *Sst*^*+*^*Chodl*^*+*^ GABAergic neurons in rare sparse cell types. In a few cases, there is variation in the mapping between independent experiments, which is consistent with biological variation, for example, co-localization of cell types (*e*.*g. Sncg* and *Vip* GABAergic neurons) is detected across slices from the same batch (*section 1* and *section 2*) but not in *section 3*; *L6 IT* cells are more localized in layer six in *slice 3*, and *Vip* neurons are more uniformly distributed in *section 3* than in *section 1* and *section 2*. Both findings are consistent with our expectations.

Importantly, Tangram correctly predicted spatial expression patterns from the mapped cells, when we withheld those genes in the training and then compared them to the Visium measurements (**Fig. 4c-f**). Specifically, we partitioned the genes into 1,237 training genes and 29,816 test genes unseen in the learning of the model, and used spatial correlation as before (**Fig. 4c**). The 90^th^ quantile of spatial correlation coefficients of training genes is >62%, setting a prediction threshold for test genes (**Fig. 4c,d**): 50% of the testing genes exceed this threshold (**Fig. 4c,d**). As the number of training genes in reduced from 1,237 to 123, so does the relative prediction accuracy (**Fig. 4d**), although it remains substantial. Inspection of spatial patterns from select test genes shows that while our predictions always result in a localized pattern in the upper layer, agreement against Visium measurements deteriorates as the gene is more sparsely detected in the original Visium experiment (**Fig. 4e**, where sparsity is defined as the fraction of voxels in which the gene is undetected).

We hypothesized that this poor agreement could be due to technical “drop-outs” (∼15,000 test genes are entirely undetected in our Visium datasets). Supporting this hypothesis, there is a strong correlation between our prediction scores and data sparsity (**Fig. 4f**): 98% of non-sparse genes (sparsity<50%) are correctly predicted by our model (spatial correlation >62% threshold, **Fig. 4f**, region i), with only ∼70 non-sparse genes that are not well predicted (**Fig. 4f**, region ii). Non-sparse test genes that are not well predicted have predicted patterns that are sparser than Visium measurements, suggesting that the disagreement might be due to drop-outs in the snRNA-seq data (**Fig. 4g**). Finally, ∼80% of the transcriptome measured in Visium is highly sparse (**Fig. 4f**, region iii; the same genes are also too low to be detected by the Allen ISH atlas). This raises the possibility that our predictions may provide more accurate estimates of the real spatial expression for such genes. Supporting this, we compared our predictions to measurement for the two genes available in both MERFISH and our sparse genes. In both cases, our predicted spatial patterns agree with MERFISH measurements (**Fig. 4h**). Future studies with additional genes could help with further validations.

### Spatial localization of chromatin accessibility patterns with SHARE-seq

We next used Tangram’s successful spatial mapping through RNA, as a scaffold to map additional molecular profiles with no available spatial data, but that can be measured by single cell multi-omics. In particular, we set to map a joint single cell RNA and ATAC-Seq, which we profiled simultaneously in the >3000 cells from whole mouse brain by SHARE-Seq^5^ (detecting ∼18,000 genes) and annotated as nine glutamatergic cell subsets (EN, excitatory neurons), five GABAergic cell subsets (IN, inhibitory neurons) and five non-neuronal subsets (A1.E1, MX, NSC, OG1, P1)^5^ (no immune cells were captured, and cortical layer subsets were not annotated). We aligned the SHARE-seq data to the MERFISH data using the snRNA-seq profiles, then transferred the snATAC-seq profiles of the cells to space, to visualize inferred spatial patterns of chromatin accessibility and transcription factor motif scores at single cells resolution (**Fig. 5**).

We mapped SHARE-seq data both probabilistically (**Fig. 5a**) and deterministically (**Fig. 5b**) and obtained cell type distributions. Our mapping reveals that EN01 are localized in layer L2/3, EN04 in layer 4, EN07 in layer 5/6, EN05 in layers 5 and 6 and EN02 in layer 6. Interestingly, IN02 seems more prominent in layer 6. Also, the non-neuronal cell type MX (labeled “Unconfirmed”^5^) appears concentrated to the bottom left part of the ROI, which does not resemble known patterns of cortical cell types. While the mapping is overall consistent, it is less biologically precise than in the previous cases, likely due to the lack of immune cells (missing “puzzle pieces” for Tangram) and the fact that the cells were not profiled specifically from the cortex.

We used the snRNA-based mapping to infer spatial patterns of chromatin accessibility and transcription factor activity (**Fig. 5c,d**), and compared them to spatial expression patterns. In some cases, gene expression is higher at a particular cortical layer but localization is not observed in the projected snATAC-seq (*C1ql3, Il1rapl2, Kcng1*). In other cases, the projected snATAC-seq forms spatial patterns, even though the corresponding predicted gene does not show a spatial pattern (*Scgn, Il4ra, Mrgprx2*). In only a minority of cases, we observed correlation between snRNA-seq and snATAC-seq patterning (*Dnase1l3, Egfr* and *Elfn1*). We similarly visualized inferred spatial patterns of transcription factor motif activity scores (identified from the snATAC-Seq profile in each cell^37^) (**Fig. 5d**). Interestingly, some of them exhibit a slightly localized pattern (*Msc, Bhlhe22, Egr2*), even for transcription factors whose predicted RNA was neither measured in MERFISH nor in SHARE-seq (e.g. *Tcfeb* and *LINE10014*).

### Integration of histological and anatomical atlas support

In the mouse brain, there are already rich histological and anatomical atlases, with careful annotations and associated data, including *in situ* hybridization of individual markers. In those cases when histological data is directly available for specimens profiled in single cell atlases, those can serve as a bridge to these pre-existing atlases. This then both helps create a full atlas, in the context of a Common Coordinate Framework (CCF) for the brain, as well as provide spatial molecular data from legacy ISH.

To this end, we developed an additional module in Tangram to connect across scales by registering histology/spatial data on an anatomically annotated CCF, such as the Allen CCF for the adult mouse brain^28^. As an alternative to methods that either require supervision or intact tissue, we combine a Siamese neural network model with a semantic segmentation algorithm to produce full segmentation masks of anatomical images (**Figs. 6** and **7**). The Siamese network model builds a latent space which allows a uniform encoding irrespective of technical artifacts in the images, such as the presence of “holes” in dissected regions (from which cells or nuclei were collected). The semantic segmentation model produces a segmentation mask with a color scheme that is compatible with the Allen ontology. Because we produce a mask with matching colors, we can then register the images automatically as we do not need to provide corresponding landmarks; instead the anatomical regions in the mask are the landmarks.

Building on methods for face recognition^16^, we learned a latent space using a Siamese network model^17^ trained on mouse brain images (**Fig. 6a, Methods**). We trained the model so that each image was encoded according to salient anatomical landmarks, whereas technical properties such as illumination or staining were factored out (**Methods**). The learned latent space displayed a one-dimensional manifold structure, where the “head” of the manifold contains images from the olfactory bulb, and the “tail”, images from cerebellum (**Fig. 6b**). The model predicted the image from the Allen CCF at the same coronal depth of our histological image. We validated the predictions by checking consistency across the whole training set (**Fig. 6c**), and by visual inspection (**Fig. 6d**). We then used the trained model to retrieve the image from the Allen CCF onto which we register our histological image.

Next, we segmented our images to generate a custom mask for our images using the same color scheme adopted by the Allen ontology. For this, we applied semantic segmentation^18^, and segmented five classes in our histological image (**Fig. 7a**): background, cortex, cerebellum, white matter and other grey matter. As the training set is scarce, we adopted a combination of transfer learning and heavy augmentation during training (**Fig. 7b**) and validated it by inspecting predictions on test atlases (**Fig. 7d**). Finally, we combined segmentation with the Siamese model, to obtain a fully automated registration pipeline (**Fig. 7c**), leveraging the fact that registering two masks (one on the Allen image and one on the image of our sample) is a simpler problem than registering the two images directly.

### A learned histological and anatomical atlas of single nuclei from the somatomotor mouse cortex

To demonstrate this final framework, we assembled an integrated atlas of the somatomotor area of the healthy adult mouse brain using publicly available atlases. As noted above, we collected ∼160,000 snRNA-seq profiles from three dissected regions of interest (ROIs) in the somatomotor area (**Fig. 8a**). We used a lab procedure^38^ where nuclei are isolated from a biopsy “punch” in a frozen dissected region. To help relate this region to the known anatomy, we also obtained stained histological sections on the punched section, which are approximately 200μm deep (**Methods**).

**Figure 8.**
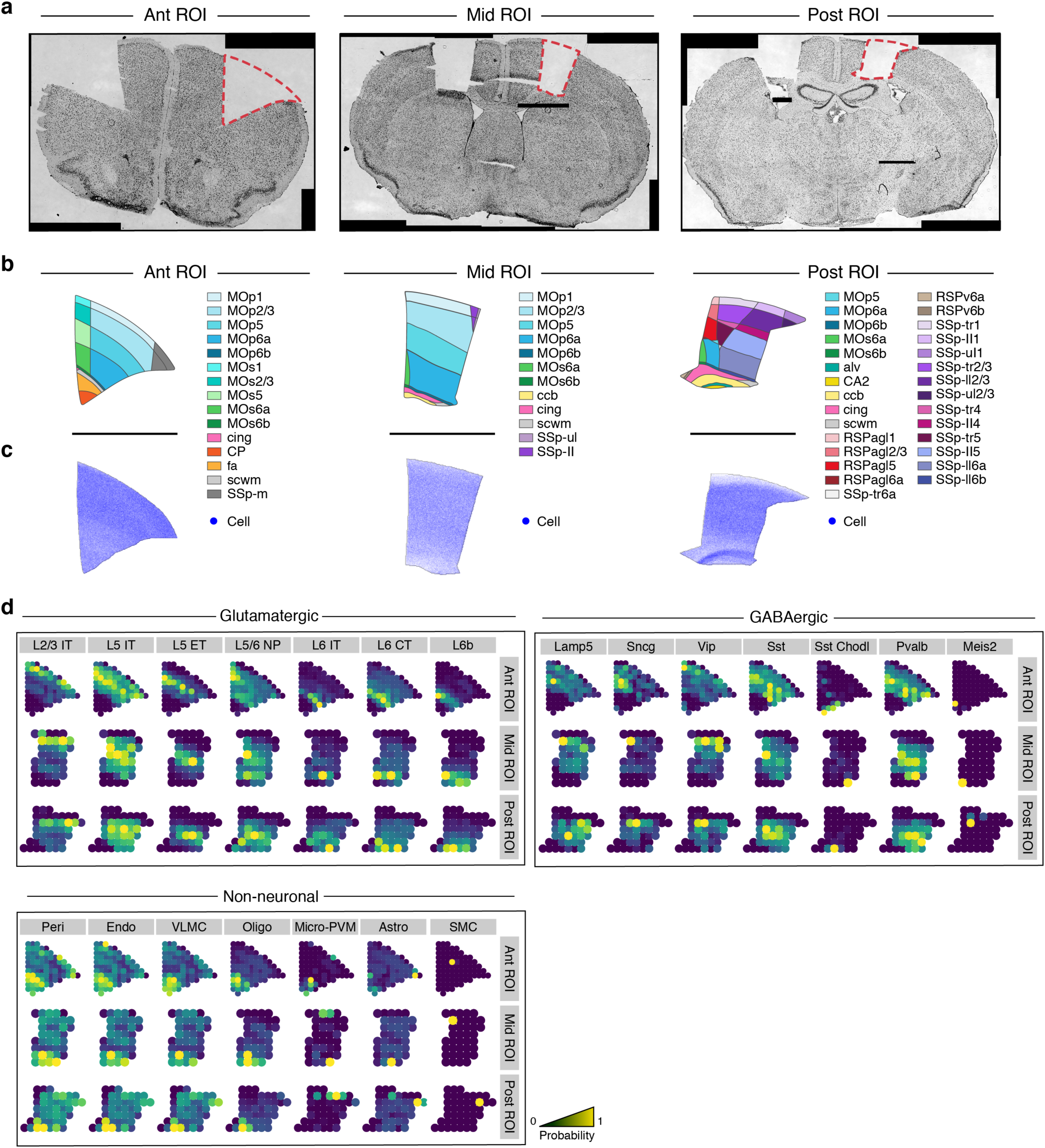
Tangram mapping of snRNA-seq profiles to histological and anatomical mouse brain atlases. **a**. Regions of interests. Nissl-stained images of coronal mouse brain slices highlighting the three regions of interest from which snRNA-seq data from the motor area were collected. **b,c**. Registration pipeline generates anatomical region and cell density maps. Anatomical region (b, color legend, from the Allen Common Coordinate Framework) and cell map (c, color bar, from the Blue Brain Cell Atlas) maps of each of the three dissected ROIs. **d**. Probabilistic mapping of snRNA-seq data on the ROI. Probability of mapping (color bar) of each cell subset (grey label) from each of 3 major categories within each ROI (rows).

We applied Tangram’s anatomical mapping module to the histological images to precisely locate the region of dissection on the Allen CCF, and then queried the *Allen Mouse Atlas* to estimate spatial gene expression at 200 *µm* resolution and the *Blue Brain Cell Atlas*^*19*^ to compute the expected cell density in each spatial voxel (**Fig. 8c**). Tangram then computed an anatomical map from the *Allen Reference Atlas*, and used it post-mapping to estimate the anatomical region to which each cell has been mapped (**Fig. 8a**). We repeated this procedure for the three ROIs, and finally mapped the snRNA-seq profiles to their corresponding ROIs. (We used the same anatomical mapping module to select nuclei for mapping from a matching histological section to that measured by Spatial Transcriptomics; above.)

The mapping predictions for cell types across the three ROIs, were self-consistent albeit less accurate than mappings using the higher resolution spatial technologies (**Fig. 8d**). Cortical layers were successfully recovered across the three ROIs, but *L5 IT* and *L5/6 NP* display a lower level of localization than in the other cases. GABAergic neurons predictions are biologically sound and we observed a more concentrated presence of *Vip+* and *Lamp5+* in upper layers, as observed with Visium-based mapping. Non-neuronal predictions did not recover sparse *mPVM* and overly concentrated *Peri, Endo* and *VLMC* cells in the subcortical part. Overall, our mapping results confirmed that glutamatergic cell type patterning is simpler to reconstruct than sparse, granular, cell type patterns typical of non-neuronal cell types, the latter requiring finer signals from modern spatial technologies.

## DISCUSSION

Gene expression in the brain and other organs exhibits a variety of spatially-organized patterns on different length scales from the microenvironment, to histology to anatomy, whose knowledge is central to unraveling biological function. Spatially resolved transcriptomic data provide an opportunity to reveal such patterns, but are currently limited by spatial resolution or the number of genes measured, and connecting them to other levels or organization can require substantial experimental efforts. Here, we harmonized snRNA-seq data with *in-situ*, histological and anatomical data, to address these limitations, towards a high resolution, integrated atlas.

Here, we developed and used Tangram to tackle various scenarios, all of which involved aligning snRNA-seq data onto a certain type of spatial data collected from the adult mouse brain cortex, from high resolution MERFISH and STARMap, through mid-resolution Spatial Transcriptomics, and to ISH associated with histological and anatomical coordinates. In each case, we validated the computational alignments by recovering consistent spatial maps of cell types and predicting the expression of holdout genes.

Each type of spatial measurement modality benefits particularly from one aspect of Tangram: for high resolution targeted datasets (smFISH, MERFISH, STARmap), Tangram expands from signature to genome-wide patterns; for lower resolution Spatial Transcriptomics dataset (Visium), Tangram yields single-cell resolution; for datasets with higher gene throughput (STARmap and Visium), Tangram detects and corrects low-accuracy gene expression patterns; for multi-modal single cell data (SHARE-seq), Tangram uses one modality to generate spatial patterns for the other, thus generating spatial multi-modal maps. Finally, histology allows registration to the Allen CCF and integration between the cellular and the anatomical scale.

In our analyses, a few hundreds of marker genes, stratified across cell types, sufficed to map the mouse brain cortex transcriptome-wide, observing consistent cell type patterns in all cases. Notably, while cell mapping can rely on even fewer genes (*i*.*e*., smFISH, 22 genes measured, **Figs. 1b,d**), we could not successfully predict transcriptome-wide spatial gene expression in that case, in contrast to our success with MERFISH (254 genes measured). This suggests that at least a few hundred marker genes are required to comprehensively map the mouse brain cortex, at least in the context of cell types. As we expand studies of the brain to other patterns – such as short-term responses that characterize more transient cell states and programs – the optimal number of marker genes required for mapping could depend not only on the number and diversity of cell types in the tissue and specificity of marker genes, but on the structure of other gene programs and their inter-relations. Tangram, using both genome-wide and targeted spatial scaffold data can help assess the extent of markers needed to capture a response.

Future applications of Tangram with more than a single spatial scaffold at a time (*e*.*g*., taking both MERFISH, Visium and snRNA-seq as inputs simultaneously) should help reconcile a complete integrated spatial atlas of gene expression in the brain. Moreover, Tangram could be applied in an iterative fashion across the modalities. For example, as we showed Tangram can correct gene expression in spatial methods when the measurements of the specific genes have lower quality. Such output can then be used as a novel spatial dataset onto which to re-align the snRNA-seq profiles. This technique, called active learning, will be explored in future studies. Finally, as additional spatial transcriptomics technologies arise, Tangram can flexibly take them as input. For example, it could be applied to SLIDE-seq2 data^39^, which has a spatial resolution of 10 μm (compared to Visium’s 50-55μm) and is more robust to technical dropouts^39^.

An interesting case study of Tangram is its ability to resolve spatial patterns of multi-modal data, when only one modality is available in the spatial scaffold, as we demonstrated using SHARE-seq data to predict spatial patterns for scATAC-seq data. This approach can be adopted with every technology that profiles the transcriptome at single cell level with an additional modality: examples include CITE-seq^40^ (cellular proteins) and Patch-seq^41^ (electrophysiological and morphological properties of neurons). Alternatively, one can independently measure various modalities in different subpopulations at single-cell level, integrate them by learning a common latent space^24,42,43^, and then input this integrated multi-modal data to Tangram to resolve multi-modal spatial patterns. Aligning multi-modal data is particularity intriguing in cases for which there is no assay for spatially resolving data of a certain modality. For example, chromatin accessibility could not be spatially resolved at single-cell level until very recently^44^.

Finally, while our work focused on a specific region in the mouse brain, Tangram is applicable to any brain region, towards its complete atlas, and to any other organ, as well as disease tissue. To integrate across scales, Tangram’s registration pipeline requires a CCF and is therefore currently applicable to a few organs. At present, the mouse brain possesses the most advanced and well-developed CCF ^28^, but efforts are underway to construct analogous reference maps for different organs^9^, towards the construction of cell atlases of all organ in mouse and human.

---

**Extended Data Figure 1.**
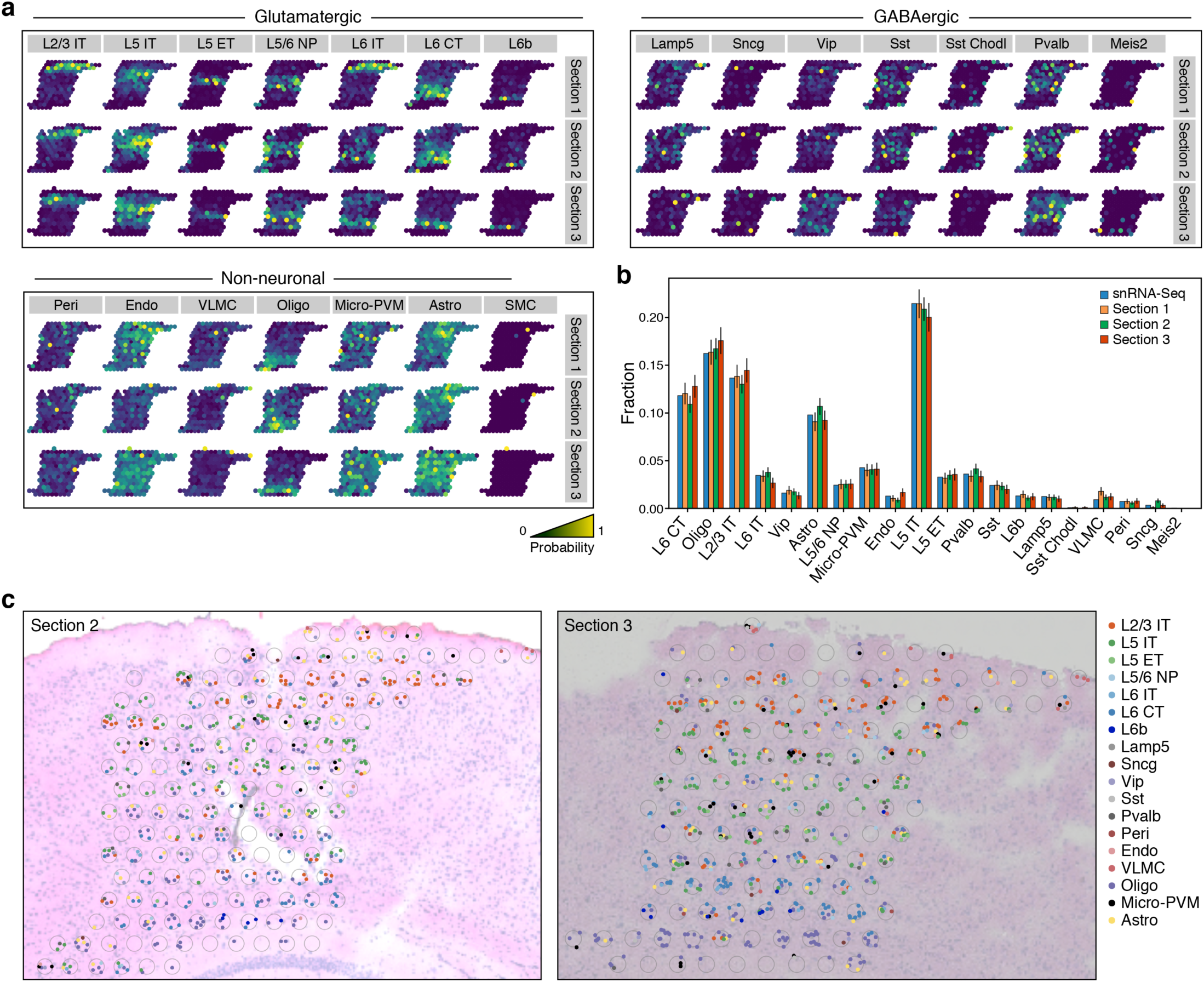
Mapping results on Visium data are consistent across three datasets. **a**. Consistent probabilistic maps across models trained from replicate datasets. Probability of mapping (color bar) of each cell subset (grey label) from each of 3 major categories in models trained separately from three Visium sections (rows). *Section I* is the same shown in **Fig. 3b. b,c**. Consistent deconvolution across models trained from replicate datasets. **b**. Fraction of cells (*y* axis) of each cell type (*x* axis) obtained after deconvolution with models trained separately by each of three Visium sections and in snRNA-seq. **c**. Predicted single cells (colored dots, legend) in each Visium voxel (grey circle) based on snRNA-seq data mapped onto Visium section 2 (left) and section 3 (right) (compare to **Fig. 3b**). Cell assignment within a voxel is random with respect to the specific segmented cell.

## Materials and Methods

### Tangram mapping algorithm

#### Introduction

We use the index *i* for cells (*i*.*e*. snRNA-seq data), *k* for genes and *j* for spatial voxels (circular spots, pucks, etc.). Our goal is to learn a spatial alignment for the cells, organized as a matrix *S* with dimensions *n*_*cells*_ × *n*_*genes*_, where *n*_*cells*_ is the number of single cells, such that *S*_*ik*_ ≥ 0 is the expression level of gene *k* in cell *i*. In order to map, we voxelize the spatial volume at the finest possible resolution (which depends on the mapping case, *e*.*g*. 200 *µm* when mapping with the Allen Brain Atlas, individual cells when mapping with MERFISH, etc.), and index the voxels in an arbitrary one-dimensional fashion. We then introduce two quantities: the *n*_*voxels*_ × *n*_*genes*_ gene expression matrix *G*, where *G*_*jk*_ ≥ 0 denotes the expression of gene *k* in voxel *j* (we do not assume that *G* and *S* measure gene expression using the same unit of measures), and a *n*_*voxel*_-long vector 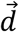 of cell densities, where 0 ≤ *d*_*j*_ ≤ 1 is the cell density in voxel *j*, and 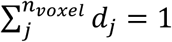.

We aim to learn a mapping matrix *M*with dimension *n*_*cells*_ × *n*_*voxels*_, such that *M*_*ij*_ ≥ 0 is the probability of cell *i* of being in voxel *j*. Therefore, we require a probability constraint 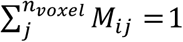. Our mapping strategy is probabilistic, perform a soft assignment. From the mapping matrix *M*, we further define two quantities: *M*^*T*^*S*, the spatial gene expression as predicted by the mapping matrix, and the vector 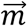 with components 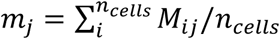 for the predicted cell density in voxel *j*. Finally, we define the softmax function along the voxel axis for any given matrix 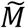 (with dimensions *n*_*cells*_ × *n*_*voxels*_). The resulting matrix *M* has elements:

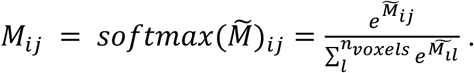

By applying the softmax, we ensure that 0 ≤ *M*_*ij*_ ≤ 1 and 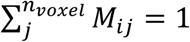.

#### Mapping algorithm

To learn the mapping matrix, we minimize the following objective function with respect to 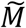 (note that in the objective we use 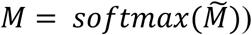:

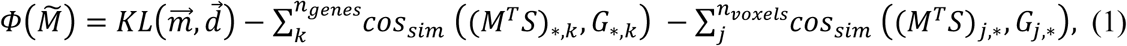

where *KL* indicates the Kullback–Leibler divergence and *cos*_*sim*_ is the cosine similarity function. The first term is the density term: we enforce that the learned density distribution is as similar as possible to the expected density. The second term is the gene/voxel expression term: it enforces that the predicted expression for each gene over the voxels is proportional to the expected gene expression over the voxels. The third term is the voxel/gene expression term: for each voxel, the predicted gene expression needs to be proportional to the expected gene expression. Minimization is obtained via gradient-based optimization using the *PyTorch* library.

Using the objective (1), Tangram maps all sc/snRNA-seq profiles onto space. If the number of sc/snRNA-seq profiles is higher than the known number of cells in the spatial data, Tangram can instead filter the sc/snRNA-seq profiles and learn the optimal subset of sc/snRNA-seq profiles that best explains the spatial data. The latter approach is explained next.

#### Mapping with a filter

We introduce a filter vector 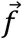 of dimension *n*_*cells*_ so that cell *i* can either be mapped (*f*_*i*_ = 1) or not mapped (*f*_*i*_ = 0). To filter, we multiply each row of the single-cell matrix, *S*_*i*,*_, and each row of mapping matrix, *M*_*i*,*_, by *f*_*i*_, as shown below in the new objective. The filter values *f*_*i*_ are learned during training, in order to retain the cells that best explain the spatial data. To explicitly promote Boolean values (*i*.*e*. 0s or 1s) in the filter values, we add a filter regularizer in the objective. To enforce the total number of filtered cells, we introduce a count term: a soft constraint in the objective that promotes a number of mapped cells in the filter equal to *n*_*target_cells*_. With this in mind, we formally define the objective. We define a real vector 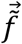 of dimension *n*_*cells*_ and define 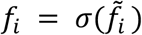, where we apply the sigmoid function *σ* to ensure that 0 ≤ *f*_*i*_ ≤ 1. We then define *S*^*f*^ = *diag*(*f*) · *S* and *M*^*f*^ = *diag*(*f*) · *M*, namely, the filtered versions of the single cell matrix and the mapping matrix, respectively. We also define 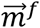, vector with components 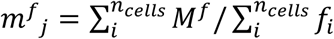, as the predicted density of filtered cells in voxel *j*. The objective function, which we minimize with respect to 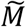 and 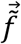, is:

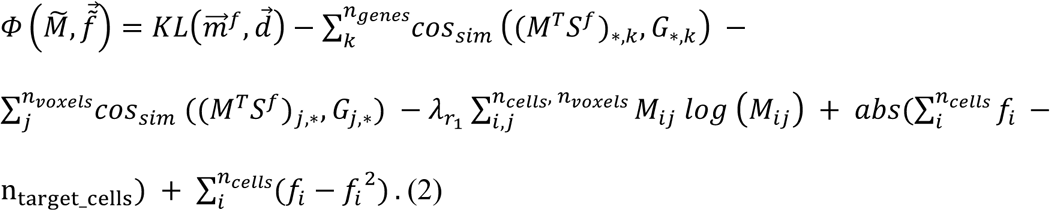

The last two terms correspond to the count term and the filter regularizer, respectively.

#### Annotations transfer

The output of the mapping algorithm is the learned mapping matrix *M* (with, optionally, the learned filter 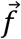). Once the mapping outcome is computed, any kind of annotation can be transferred from the sc/snRNA-seq data onto space.

We define *A* as the annotation matrix with dimensions *n*_*cells*_ × *n*_*annotations*_, where *n*_*annotations*_ is the number of different annotations characterizing single cells (*e*.*g*., genes, cell types, or any other modality). If annotations are categorical values (such as cell types), we one-hot encode them over multiple columns in *A*. Annotations in space are retrieved by computing:

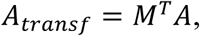

or, if the filter has also been learned, via:

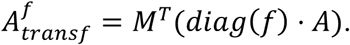

The computed matrix *A*_*transf*_ has dimensions *n*_*voxel*_ × *n*_*annotations*_, and therefore denotes the annotations in space.

#### Cell type calling

When *A* describes cell types, *A*_*transf*_ describes the probabilistic counts for each cell type in each cell voxel. This corresponds to *probabilistic mapping* and can be interpreted as the mixture of cell types which best explain the *in situ* gene expression. When the technology allows for single-cell spatial resolution, voxels correspond to single cells in space. In this case, probabilistic mapping can be seen as an unnormalized probability distribution over cell types for the voxel/cell. As a consequence, for technologies with single-cell spatial resolution, we can define a *deterministic mapping* as the mapping assigning the *most likely* cell type in this distribution to each spatial voxel/cell.

#### Mapping spatial data from targeted technologies

smFISH (**Fig. 1**), MERFISH (**Fig. 2**), and STARMap (**Fig. 3**) allow for single-cell spatial resolution, therefore, the number of spatial voxels needs to be equal to the number of cells. As snRNA-seq profiles are typically more numerous than MERFISH voxels, we can use the mapping with the learned filter, namely, Eq. (2). In this case, *n*_*target_cells*_ = *n*_*voxel*_ and the expected density 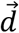 is uniform and equal to 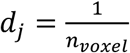 for all *j*. This enforces that each cell is mapped to one voxel only and vice versa. If the number of available single cells is lower than the number of spatial spots, we can instead map with Eq. (1), using the same constant density 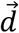.

For the MERFISH case, we mapped 58,022 10Xv3 snRNA-seq profiles in 4,951 spatial spots. From the 26,944 genes in the snRNA-seq data, we selected 1,408 marker gene as the top 100 marker genes stratified across the 22 cell types. We mapped using the intersection between these marker genes and the 254 MERFISH genes, which corresponded to 253 genes.

For the smFISH case, we mapped 11,759 SMART-Seq2 snRNA-seq profiles in 4,840 spatial spots. In this case, 40,056 transcripts are measured in the snRNA-seq data. Only 22 genes were measured in smFISH, all of which were also present in the snRNA-seq data. Therefore, we used all 22 genes for mapping.

For the STARMap case, we used the same snRNA-seq data as for smFISH, which we mapped on 1,550 spatial spots. We took the intersection of 995 genes between the 1,020 STARmap transcripts and the 40,056 transcripts in the snRNA-seq data. We used these 995 genes for mapping. The algorithm converges after 1,200 epochs in all the experiments.

#### Mapping Visium data

We began by identifying the Post ROI on the Visium histological image using our registration pipeline (below). Next, we segmented the cells within the ROI using the software ilastik (https://www.ilastik.org). We then built the density vector 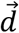, by computing the cell density inside each voxel (*i*.*e*., Visium circle, as in **Fig. 1e**). We mapped using the objective described in Eq. (1). Mapping yields the matrix *M*, which we used to generate probability maps for the cell types within the ROI. To deconvolve, we mapped using Eq. (2), to constrain the expected number of cells in each Visium voxel. Specifically, we used *n*_*target_cells*_ = *n*_*seg*_, where *n*_*seg*_ is the total number of segmented cells in the Visium ROI, to enforce that only a subset of cells is actually mapped. The count term combined with the density term led to the expected number of mapped cells in each Visium voxel. After training, we assigned the types of the cells mapped within each voxel randomly to specific segmented cells within that voxel.

For probabilistic mapping on Visium data, we ran the optimizer for 300 epochs to reach convergence. At the end, more than 99% cells were assigned to an individual voxel with probability greater than 50%. For used deterministic mapping in deconvolution, we trained the optimizer for 6,000 epochs to reach convergence. At the end, more than 99% cells were assigned to an individual voxel with probability greater than 50%. For *section 1* dataset, the number of cells filtered (*f*_*i*_ > 0.5) is 880 (89% of segmented cells). Segmented cells for which there is no filtered mapped cell are not shown in the figures.

For both probabilistic and deterministic mapping, we used 58,022 10Xv3 snRNA-seq profiles for 162, 161 and 134 spatial spots, respectively, in *section 1, section 2* and *section 3*. Among the 26,944 transcripts in the snRNA-seq data, 1,408 marker genes were selected. We mapped using the intersection of these genes with Visium genes (31,053), corresponding to 1408 genes.

#### Mapping Allen atlas data

We used 58,022 10Xv3 snRNA-seq data for 83, 38 and 43 spatial spots, respectively, in the *anterior, mid* and *posterior* ROIs. Among 26,944 transcripts in the snRNA-seq data, 1,408 marker genes were selected. We mapped using the intersection between these genes with Allen atlas genes measured coronally (overall, 4,345 genes); the intersection corresponds to 750 genes. The algorithm converged after 150 epochs.

### Data collection - snRNA-seq data and histological images

#### Mouse experiments

Mice were group housed with a 12-hour light-dark schedule and allowed to acclimate to their housing environment for two weeks post arrival. All procedures involving animals at MIT were conducted in accordance with the US National Institutes of Health Guide for the Care and Use of Laboratory Animals under protocol number 1115-111-18 and approved by the Massachusetts Institute of Technology Committee on Animal Care. All procedures involving animals at the Broad Institute were conducted in accordance with the US National Institutes of Health Guide for the Care and Use of Laboratory Animals under protocol number 0120-09-16.

#### Brain preparation prior to anatomical dissection and snRNA-seq

At 60 days of age, C57BL/6J mice (50% males, 50% females) were anesthetized by administration of isoflurane in a gas chamber flowing 3% isoflurane for 1 minute. Anesthesia was confirmed by checking for a negative tail pinch response. Animals were moved to a dissection tray and anesthesia was prolonged via a nose cone flowing 3% isoflurane for the duration of the procedure. Transcardial perfusions were performed with ice cold pH 7.4 HEPES buffer containing 110 mM NaCl, 10 mM HEPES, 25 mM glucose, 75 mM sucrose, 7.5 mM MgCl_2_, and 2.5 mM KCl to remove blood from the brain and other organs sampled. The brain was removed immediately and frozen for 3 minutes in liquid nitrogen vapor and moved to −80°C for long term storage. A detailed protocol is available at protocols.io (dx.doi.org/10.17504/protocols.io.bcbrism6).

#### Generation of MOp dissectates and snRNA-seq

Frozen mouse brains were securely mounted by the cerebellum onto cryostat chucks with OCT embedding compound such that the entire anterior half, including the primary motor cortex (MOp), was left exposed and thermally unperturbed. Dissection of 3 consecutive 500μm anterior-posterior (A-P) spans of the MOp was performed by hand in the cryostat using an ophthalmic microscalpel (Feather safety Razor #P-715) precooled to −20°C and donning 4x surgical loupes. Each 500μm step was accomplished by advancing the cryostat (Leica CM3050S) 100μm 5 times in trimming mode and cutting out each dissectate 100μm at a time. This stepwise approach serves to ameliorate disruption of the brain tissue surface that occurs with large steps. Each excised tissue dissectate pool was placed into a pre-cooled 0.25 ml PCR tube using pre-cooled forceps and stored at −80°C. In order to assess dissection accuracy, 10μm coronal registration sections were taken at each of the 500μm A-P dissection junctions and imaged following Nissl staining. Nuclei were extracted from the frozen tissue dissectates using gentle, detergent-based dissociation, according to a protocol (<u>dx.doi.org/10.17504/protocols.io.bbseinbe</u>) adapted from one generously provided by the McCarroll lab, and loaded into the 10x Chromium V3 system (10x Genomics). Reverse transcription and library generation were performed according to the manufacturer’s protocol.

snRNA-seq data were analyzed using the scanpy package^45^. snRNA-seq data were pre-processed via the following steps: we removed cells with high mitochondrial gene content; we normalized the data to correct for library-size; we applied the function *f*(*x*) = log (1 + *x*) to the normalized counts. The resulting snRNA-seq data are ready to be mapped with Tangram. To compute marker genes of snRNA-seq data, we applied a computational pipeline described in the tutorial of the scanpy package available at https://scanpy-tutorials.readthedocs.io/en/latest/pbmc3k.html.

We used normalized quantities to visualize gene expression via mRNA counts (**Figs 1b, 1c, 1f, 2c, 2e, 2f, 3c, 3d, 3e, 4e, 4g, 4h, 5c** and **5d**), gene expression via fluorescence (**Figs. 2c, 2f, 3c, 4h**, and **5c**) chromatin accessibility via ATAC peak counts (**Figs. 1f** and **1c**), and transcription factor activity via z-scores (**Fig. 5d**). Normalization is performed by rescaling the colorbar in each image, so that the minimum (respectively, maximum) value of the image correspond to the color with minimum (respectively, maximum) value in the colorbar. This is the default behavior of the plotting functions of the Python library matplotlib (https://matplotlib.org) which we used throughout the manuscript.

### Data collection - Visium

#### Mice

All mouse work was performed in accordance with the Institutional Animal Care and Use Committees (IACUC) and relevant guidelines at the Broad Institute, with protocol IACUC 0147-02-17.

#### Tissue processing

Fresh-frozen wild type C57BL/6 whole mouse brain was embedded in OCT (TissueTek Sakura) and cryo sectioned at 10 µm thickness at −20°C. Tissue sections were placed in 6.5 mm squared capture areas on pre-cold Visium Tissue Optimization slides (3000394, 10x Genomics) and Visium Spatial Gene Expression slides (2000233, 10x Genomics) and adhered by warming the backside of the slides and placed at −80°C for up to 3 days.

#### Visium spatial gene expression library generation

The tissue optimization sample slide and spatial gene expression slide were processed according to the manufacturer’s protocols (CG000238_VisiumSpatialTissueOptimizationUserGuide_Rev_A.pdf and CG000239_VisiumSpatialGeneExpression_UserGuide_Rev_A.pdf). Briefly, tissue sections were warmed to 37°C for 1 minute and fixed for 30 min in ice-cold methanol, followed by 1 min isopropanol incubation at room temperature. Tissues were then hematoxylin and eosin (H&E) stained according to the protocol. Morphology brightfield images were taken with a Zeiss Axio microscope with the Metafer slide-scanning platform (Metasystems) with a 10x objective. For the tissue optimization slide fluorescent images, a TRITC filter and 10x resolution was used. Images were joined together with the VSlide software (Metasystems) and exported as tiff files. To optimize tissue permeabilization time, six different time points with 3-minute increments were tested on the tissue optimization sample slide. 12 minutes of permeabilization was used for the spatial gene expression slide. RNA released from the tissue was converted to cDNA by priming to the spatial barcoded primers on the glass via reverse transcription in the presence of template switching oligo, to generate full-length, spatially barcoded, UMI containing cDNA. Subsequently, following second strand synthesis, a denaturation step released the cDNA, followed by PCR amplification. Finally, sequencing-ready, indexed spatial gene expression libraries were constructed. Two of the libraries were pooled together and sequenced on a NextSeq 500/550 High output kit at 1.8 pM concentration. The sequencing settings were: read 1, 28 cycles; read 2, 91 cycles; index 1, 8 cycles.

#### MOp Visium raw read processing

Raw FASTQ files generated by Illumina’s BCL2FASTQ conversion and the histology H&E images were provided as input to the SpaceRanger software (10x Genomics) version 1.1.0, available at https://support.10xgenomics.com/spatial-gene-expression/software/downloads/latest. Sequencing reads were mapped to the mm10 reference mouse genome using STARv2.5 mapping as part of SpaceRanger suite. Spatial barcodes were assigned by SpaceRanger to the barcoded spatial spots and aligned with the tissue image with the aid of the fiducial frames. Barcodes/UMI and genes were counted for the individual spots to generate an output gene expression per-spot matrix used as input for downstream data analysis.

#### MOp MERFISH data preprocessing

We obtained MERFISH data from the Zhuang lab. We preprocessed the data to remove sub-cortical cells. To identify sub-cortical cells, we identify cells overly expressing *Nxph4* (a marker gene of L6b region) and fit those cells with a square root polynomial. All cells below the fit were considered sub-cortical and removed.

### Image datasets for registration pipeline

To locate ROIs, we used images of Nissl-stained coronal mouse brain slices collected in the Macosko lab. To train and test the models presented in **Fig. 6-8**, we used the following public image datasets:

- (dataset **avg**): 1,320 images/segmentation masks of coronal slices from the average template of the Allen adult mouse brain atlas at resolution 10μm (available at http://download.alleninstitute.org/informatics-archive/currentrelease/mouse_ccf/average_template/slice_images/).
- (dataset **ara**): 1,320 images/segmentation masks of coronal slices from the Nissl template of the Allen adult mouse brain atlas at resolution 10μm (available at http://download.alleninstitute.org/informatics-archive/current-release/mouse_ccf/ara_nissl/).
- (dataset **p56c**): 132 images/segmentation masks of coronal slices from the Allen P56 coronal reference atlas (<u>available at</u> https://mouse.brain-map.org/experiment/thumbnails/100048576?image_type=atlas).
- (dataset **p56d**): 504 images of coronal slices from the Allen Development Atlas P56 (<u>available at</u> http://help.brain-map.org/display/atlasviewer/Allen+Developing+Mouse+Brain+Atlas).
- (dataset **brainmaps**): 111 images of coronal slices from Nissl-stained BrainMaps atlas (ID: 43) (available at http://brainmaps.org/index.php?action=viewslides&datid=43), and 87 images of coronal slices from Nissl-stained BrainMaps atlas (ID: 38) (available at http://brainmaps.org/index.php?action=viewslides&datid=38).
- (dataset **ish**): 30 images of coronal slices from the Allen ISH Data (available at. https://mouse.brain-map.org/search/index).

#### Siamese network model for depth calling

We used datasets **avg, ara, p56c** and **p56d** for training. Training images were resized to 224×224 and casted to type float32. Pixel values were rescaled between zero and one, prior to training. All images were augmented using <u>imgaug</u> (https://github.com/imgaug) library. As training labels we used numerical coordinates, indicating the spatial coronal depth (*i*.*e*., posterior) of each mouse brain image on a scale of 10 *µm*. For the **avg** and **ara** datasets, labels were readily available from their tensor coordinates. Labels for the **p56c** and **p56d** datasets were also readily obtained using the *AllenSDK API* (https://allensdk.readthedocs.io/en/latest/). Dataset **brainmaps** and **ish** were manually annotated and used as test sets.

In designing the Siamese network model, we used a DenseNet169 encoder pretrained on the ImageNet dataset and open-sourced through Keras Applications. We fine-tuned the encoder by training the last convolutional layer. We added two fully connected layers on top of the encoder in order to map the extracted features to our 512-dimensional latent space. A last fully-connected layer was used to map the latent space to the model output as represented in **Fig. 6**. All fully connected layers were trained.

A training sample consisted of two random images from the annotated datasets. The difference in spatial depth coordinates between the two images, denotes by 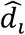, was used as the label. For example, if the first image were at posterior (depth) 500 *µm* and the second at a posterior 700 *µm* the corresponding label would be 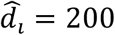. We used as penalty the mean squared error between the spatial depth difference predicted by our network *d*_*i*_, and the labels 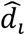:

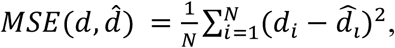

where *N* indicates the number of training samples. We trained the model for 50 epochs using 18,000 image pairs per epoch, partitioned to batches of 16 images.

#### Semantic segmentation model for anatomical region calling

We used datasets **avg, ara** and **p56c** as training sets, since masks were available. Training images were resized to 512×512 and casted to type float32. Pixel values were rescaled between zero and one. As labels, we used superimposable segmentation masks with the same dimension as the training images. Each mask was one-hot encoded into a 5-channel tensor to annotate each pixel into five different classes (**Fig. 7**): background (black), cortex (green), cerebellum (yellow), other grey matter (grey), and white matter (brown). We used colors consistent with the Allen ontology to facilitate registration. For **avg** and **ara** datasets, we used masks from the Allen CCFv3 ontology 2017 (available at http://download.alleninstitute.org/informatics-archive/current-release/mouse_ccf/annotation/ccf_2017/annotation_10.nrrd). For the **p56c** dataset, we downloaded the SVG masks from the Allen Institute website, and rendered them into images using the library released in this study, which builds on <u>Cairo SVG</u> (https://cairosvg.org). Both images and masks were augmented using the same pipeline adopted for the Siamese model. In transforming the masks, we ensured that the one-hot structure was preserved in the masks after augmentation.

We used a semantic segmentation model from the Tensorflow Keras version of the <u>segmentation_models</u> library (https://github.com/qubvel/segmentation_models). Specifically, we chose a U-NET architecture^18^ with a ResNet50 backbone^26^. All weights have been randomly initialized following the He scheme, with the exception of the ResNet50 encoder which was pre-trained on ImageNet. The model was trained to optimize the superposition of the cross entropy and Jaccard index (*i*.*e*. intersection-over-union). The loss function is defined as:

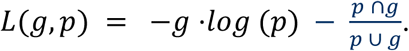

Where *g*a is the ground truth image and *p* is the corresponding model prediction. The model last unit employs a softmax activation function, thus outputting the probability of each pixel to be in each of the five classes. By applying an argmax function, we assign each pixel to its most probable class. Finally, we relied on test-time augmentation to increase model performances: each test image was augmented twelve times, and final predictions were de-augmented and averaged.

## Code availability

Tangram code is available at https://github.com/broadinstitute/Tangram, along with the datasets used to generate Figure 1.

## Author contribution

TB and AR conceived and designed the study. GS and TB developed the mapping algorithm. LB and TB developed the computer vision pipeline. RA, ZL, AS, NT, LB, GS and TB analyzed the data. CV and EM generated the MOp snRNA-seq dataset. MZ and XZ generated the MERFISH dataset. SM generated the SHARE-seq dataset, with guidance by AR and JB. AS and IAD generated *section 1* and *section 2* of the MOp Visium dataset. AR, TB, GS and LB wrote the manuscript, with input and comments from all authors.

## Acknowledgments

We thank Leslie Gaffney for her help with figure preparation and Anna Hupalowska for help with graphics. TB acknowledges useful discussions with Ed Lein, Graham Heimberg and Gokcen Eraslan. We thank all members of the Regev lab for their support. Work was supported by the BICCN grant 1U19 MH114821 (Huang and Arlotta), the Human Biomolecular Atlas Project (NIH 1OT2OD026673-01), the Klarman Cell Observatory and HHMI (AR). This publication is part of the Human Cell Atlas - www.humancellatlas.org/publications.

